# Pleiotropic Roles for the *Plasmodium berghei* RNA Binding Protein UIS12 in Transmission and Oocyst Maturation

**DOI:** 10.1101/2020.12.21.423818

**Authors:** Katja Müller, Olivier Silvie, Hans-Joachim Mollenkopf, Kai Matuschewski

## Abstract

Colonization of the mosquito host by *Plasmodium* parasites is achieved by sexually differentiated gametocytes. Gametocytogenesis, gamete formation and fertilization are tightly regulated processes, and translational repression is a major regulatory mechanism for stage conversion. Here, we present a characterization of a *Plasmodium berghei* RNA binding protein, UIS12, that contains two conserved eukaryotic RNA recognition motifs (RRM). Targeted gene deletion resulted in viable parasites that replicate normally during blood infection, but form fewer gametocytes. Upon transmission to *Anopheles stephensi* mosquitoes, both numbers and size of midgut-associated oocysts were reduced and their development stopped at an early time point. As a consequence, no salivary gland sporozoites were formed indicative of a complete life cycle arrest in the mosquito vector. Comparative transcript profiling in mutant and wild-type infected red blood cells revealed a decrease in transcript abundance of mRNAs coding for signature gamete-, ookinete- and oocyst-specific proteins in *uis12(-)* parasites. Together, our findings indicate multiple roles for UIS12 in regulation of gene expression after blood infection in good agreement with the pleiotropic defects that terminate successful sporogony and onward transmission to a new vertebrate host.

## Introduction

During the complex life cycle of *Plasmodium*, malarial parasites switch between different host cells, extra- and intracellular life cycle forms, and an invertebrate and a vertebrate host. As the parasites progress in their developmental program, and particularly after switching host, adaptation to new environmental conditions, including temperature and immune defense, is central. For instance, *Plasmodium* gametocytes must remain infectious in the blood of the vertebrate host until they are eventually transmitted to the vector during a blood meal. The gametocytes need to quickly adapt to new environmental factors, leave the digestive environment of the blood meal, and colonize the mosquito midgut (Billker et al., 2004; Billker et al., 1998). To swiftly accomplish a host switch, transcription and translation are tightly coordinated.

Transcriptional reprogramming is controlled by the apicomplexan Apetala-2 (Api-AP2) transcription factors (Balaji et al., 2005; Painter et al., 2011). Many genes needed for gametocytogenesis and early events of mosquito colonization are transcribed by the interplay of the *Plasmodium* transcription factors AP2-G and AP2-I, while asexual gene expression is repressed by AP2-G2 and AP2-FG controls female-specific gene regulation (Josling et al., 2020; Sinha et al., 2014; Yuda et al., 2015; Yuda et al., 2020). This transcriptional switch leads to the expression of mRNAs which are required for the multiple steps from gametocytogenesis to zygote formation until the motile ookinete stage is reached and the next checkpoint in transcription control is initiated by the AP2-O 1-4 family (Kaneko et al., 2015; Modrzynska et al., 2017). Therefore, a tight fine-tuning is essential to synthesize tailored proteins at the right time.

Translational repression occurs during host switch to keep gametocytes and sporozoites in a latent state. Accordingly, translation of pre-synthesized mRNAs, which are required for mosquito colonization, occurs only after transmission and constitutes a mechanism to save energy. In gametocytes a large number of genes have been described to be translationally repressed (Lasonder et al., 2008; Lasonder et al., 2016; Le Roch et al., 2004; Mair et al., 2006; Mair et al., 2010; Zhang et al., 2010). Translational repression is particularly dominant in female gametocytes and affects a wide range of mRNAs, including those that encode surface proteins, *e*.*g*. P25, P28, and CCP2, transcriptional regulators, *e*.*g*. AP2-O, and proteases, *e*.*g*. plasmepsin IV. A conserved U-rich 47 nucleotide-long cis-acting consensus motif in the 5’ and 3’UTRs of a subset of translationally repressed transcripts in *P. berghei* gametocytes was shown to be important in translational regulation in gametocytes (Braks et al., 2008; Hall et al., 2005). In the female gametocyte the DDX6-class DEAD box RNA-helicase DOZI (development of zygote impaired) is bound to its target transcripts in a large storage mRNP complex and silences their translation (Guerreiro et al., 2014; Mair et al., 2006; Mair et al., 2010). The Pumilio-family RNA binding protein Puf2 increases proliferation and decreases stage differentiation. As a consequence, deletion of *P. berghei* and *P. falciparum Puf2* resulted in a higher gametocyte rates and premature formation of exo-erythrocytic forms (Miao et al., 2010; Gomes-Santos et al., 2011; Müller et al., 2011; Lindner et al., 2013). In *pfpuf2(-)* parasites an up-regulation of many gametocyte specific transcripts, mostly independent of those that are regulated in *dozi(-)/ cith(-)*, was observed (Miao et al. 2010). Specific binding of *pfs25, pfs28* and *Plasmepsin VI* mRNA by the *Pf*Puf2 protein was shown by RNA immunoprecipitation (Miao et al., 2013).

It is likely that additional factors participate in the regulation of post-transcriptional control of gene expression during host switch to the mosquito vector. In the present study we characterized an RNA binding protein, originally identified as up-regulated in infectious sporozoites gene 12 (*UIS12*), which is also highly expressed in gametocytes. A suppression subtractive hybridization screen aimed at discovering genes which are up-regulated in salivary gland sporozoites first described the *P*. *berghei* RNA binding protein UIS12 (PBANKA_0506200) (Matuschewski et al., 2002). As expected, protein prediction programs indicate an intracellular localization. Comparison of transcriptome and proteome data indicate that *P. berghei UIS12* is expressed in sporozoites, but the corresponding protein is not detectable, suggesting that UIS12 expression is regulated post-transcriptionally (Lasonder et al., 2008; Le Roch et al., 2004; Lindner et al., 2019). Employing experimental genetics, the role of *UIS12* for parasite life cycle progression was analyzed with an emphasis on a potential function in post-transcriptional regulation.

## Results

### The RNA binding protein PbUIS12 is expressed at multiple life cycle stages

We initiated our analysis by an NCBI conserved domain database (CDD) search (Marchler-Bauer et al., 2015), which revealed two ∼70 amino acid long RNA-recognition motifs (RRM), belonging to the class of RNA-binding domains (RBD) (Dreyfuss et al., 1988), in the *P. berghei* UIS12 protein sequence. The *UIS12* gene, and particularly the two RRM domains, are highly conserved between different *Plasmodium* UIS12 orthologs (Figure 1 A), while in a candidate *Toxoplasma gondii* ortholog (TGME49_268380) the conservation is limited to the two RRM domains only. In general, the amino- and carboxy-terminal regions are less conserved, and all *Plasmodium* ortholog proteins have similar sizes of ∼1,300 amino acid residues. The *PbUIS12* coding region has a total length of 4,070 base pairs and consists of two exons (Figure 2 A). Of note, the two regions encoding the RRM domains are separated by the single intron, which also exhibits a degree of conservation.

**Figure 1:**
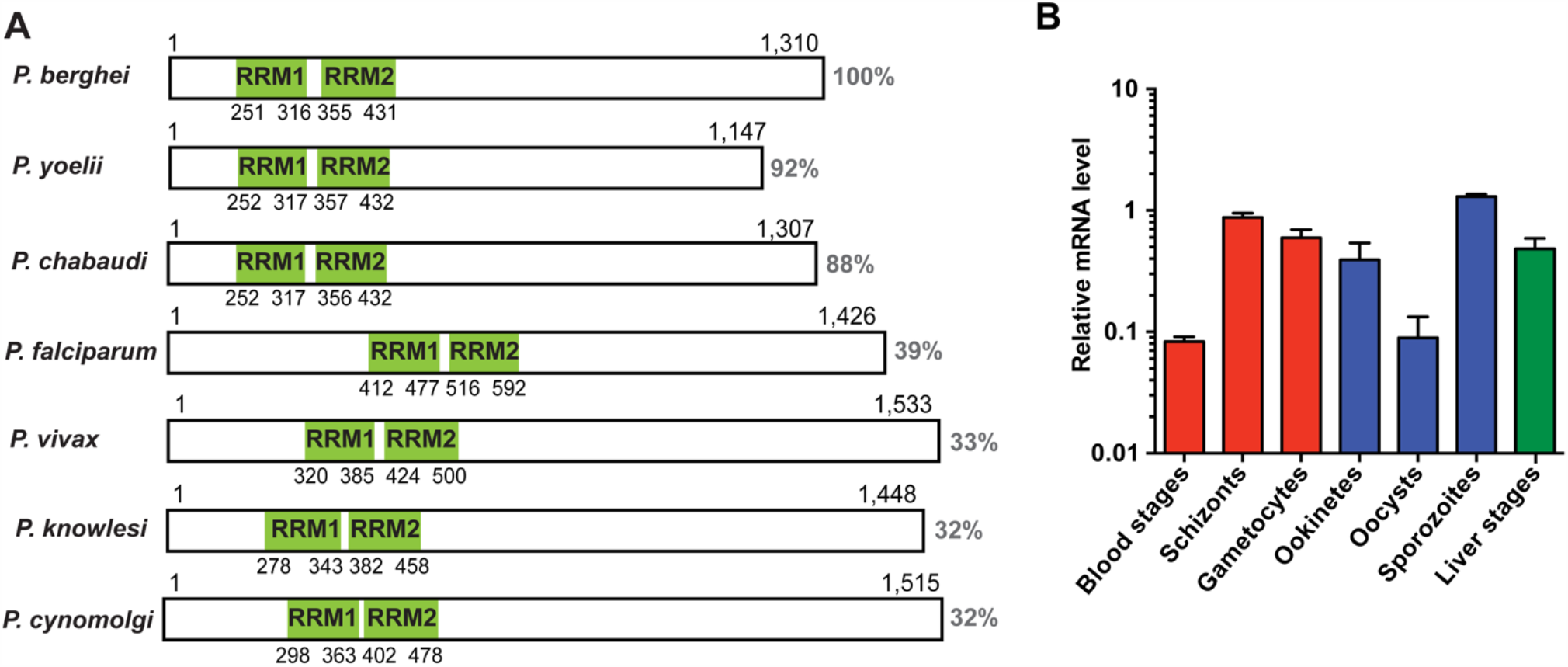
Expression analysis of UIS12. (A) Schematic display of protein primary structure illustrating length, homology and position of the two predicted RRM domains (green) of the UIS12 orthologues of different *Plasmodium* species. Protein sequence identity is shown to the right. Gene identification and accession numbers are as follows: *P. berghei*, PBANKA_0506200; *P. yoelii*, PY17X_0507300; *P. chabaudi*, PCHAS_0506300; P. *falciparum*, PF3D7_0823200; *P. vivax*, PVX_001965; *P. knowlesi*, PKNH_0606300; *P. cynomolgi*, PCYB_061630. (B) Expression profile of *UIS12* in different *P. berghei* life cycle stages; mixed blood stages, schizonts, gametocytes (red), ookinetes, oocysts, sporozoites (blue) and 24h liver stages (green). Expression levels were quantified by qRT-PCR, and data were normalized to GFP, constitutively expressed under the *PbEF-1α* promoter (cl507) (Janse et al., 2006a). Mean values (± SD) from two independent experiments shown.

**Figure 2:**
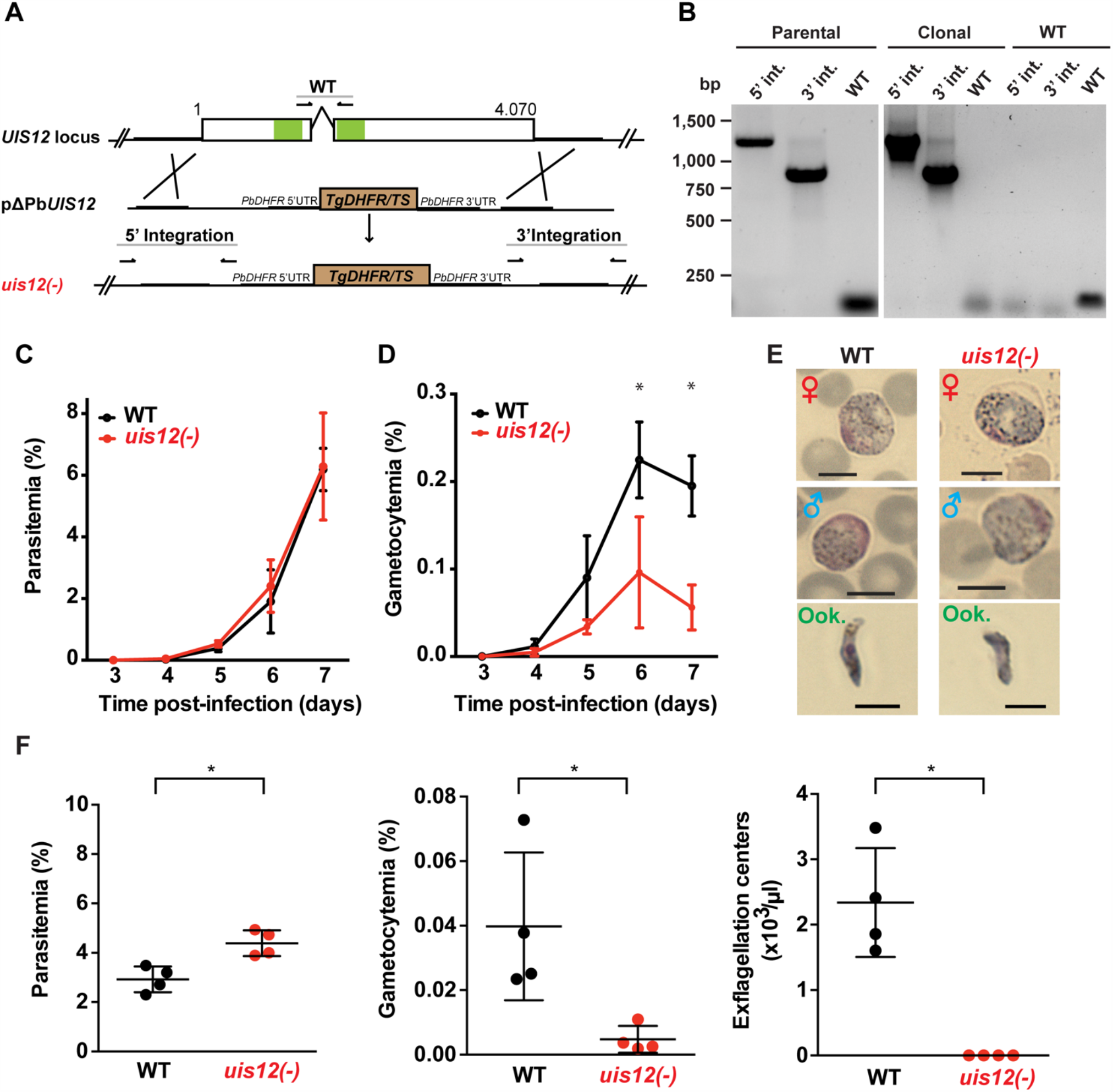
Normal asexual blood stage growth in *uis12(-)* parasites, but reduced gametocyte development and exflagellation. (A) Graphical scheme showing the replacement strategy to generate *uis12(-)* parasites, and *UIS12* gene structure. The *UIS12* locus was replaced by double-homologous recombination with the linearized replacement plasmid pΔPBANKA_UIS12, which includes a resistance marker (*Toxoplasma gondii DHFR/TS*) and the 5’ and 3’ flanking regions of *UIS12*. Arrows and lines indicate positions of knockout- and wild-type-specific oligonucleotide positions and PCR fragments, respectively. The RNA recognition motif domains (RRM) are shown in green. (B) Confirmation of *UIS12* gene disruption by diagnostic PCR. Genomic DNA of parental *uis12(-)*, clonal *uis12(-)* clone 2 and wild-type parasites served as templates. 5’- and 3’-integration-specific primer combinations amplify the predicted fragment only in the recombinant locus. Wild-type-specific primer pairs that do not produce a PCR fragment in the recombinant locus amplified the wild-type locus (right panel). Absence of residual wild-type confirms a clonal *uis12(-)* line (central panel). Time courses of parasitemia (C) and gametocytemia (D) of *uis12(-)* (clone 1, red) and wild-type (WT) (black) parasites, starting three days after intravenous injection of 10,000 mixed blood stages into C57BL/6 mice (*n*=5). Daily microscopic analysis of Giemsa-stained blood films was used to determine parasitemia and gametocytemia. (C) Parasitemia, percentage of asexual parasites per total red blood cells (D) gametocytemia, percentage of gametocytes per total erythrocytes. Means of parasitemia and gametocytemia (±SD) are shown. *, p<0.01 (multiple t-tests, one per row). (E) Gametocytes and ookinetes of *uis12(-)* (clone 1) are morphologically indistinguishable from wild-type (WT). Shown are microscopic images of Giemsa-stained female (♀) and male (♂) gametocytes and ookinetes (Ook., from ookinete cultures). Brightfield microscopy, magnification 1,000-fold. Scale bars, 5 µm. (F) Asexual and sexual blood stage development. Infected blood (C57BL/6 mice; *n*=4) was analyzed four days after injection of 10^7^ *uis12(-)* (clone 1, red) or WT (black)-infected erythrocytes, respectively. Plots shows parasitemia (left), gametocytemia (center) and numbers of exflagellation centers per microliter of mouse tail blood (right). Results are expressed as means (± SD). *, p<0.1 (Mann-Whitney test).

We next measured *UIS12* transcript abundance throughout the *P. berghei* life cycle by quantitative RT-PCR (Figure 1B). In good agreement with the previous notion of up-regulation in salivary gland sporozoites, *UIS12* mRNA steady state levels are low in midgut oocysts and highest in salivary gland sporozoites. High *UIS12* transcript levels were also detected in early (24h) liver stages. In mixed asexual blood stages *UIS12* mRNA levels are low, whereas in schizonts, gametocytes and ookinetes, considerable expression was detected. Our observations mostly corroborate published data (Howick et al., 2019; Otto et al., 2014). However, we did not specifically analyze ring stages or distinguished between male and female gametocytes (Yeoh et al., 2017). Together, *UIS12* is expressed at multiple points of the *P. berghei* life cycle, indicative of multiple functions during life cycle progression and parasite stage conversion.

### Targeted gene disruption of the RNA-binding protein *PbUIS12*

In order to study the gene function, we generated a *UIS12* loss-of-function mutant. *PbUIS12* deletion was generated by double homologous recombination employing a replacement strategy, as described previously (van Dijk et al., 1995) (Figure 2 A).

The *UIS12* targeting construct (pΔPBANKA_UIS12) comprised ∼500 bp each of the 5’ and 3’ non-coding sequences flanking a *TgDHFR*/*TS* pyrimethamine resistance cassette for positive selection with the anti-malarial drug pyrimethamine. Upon a double homologous recombination event, the *UIS12* gene is expected to be replaced by the selectable marker (Figure 2 A). As recipient parasites we used *P. berghei* (strain: ANKA; clone: cl507), which expresses the fluorescent protein GFP under the control of the constitutive *eIF1α* promoter (Janse et al., 2006a). After transfection we recovered parental *uis12(-)* blood stage populations, from which clonal lines were generated by injection of limiting serial dilutions into mice. The desired recombination event was confirmed by diagnostic PCR, which confirmed the presence of 5’ and 3’ integration in the parental and the clonal *uis12(-)* populations, and by the absence of wild-type-specific PCR fragments in the clonal *uis12(-)* populations. During the course of our *UIS12* knock-out analysis and to further corroborate our findings we successfully generated an independent second *uis12(-)* knockout line (*uis12(-)* clone 2), in a different *P. berghei* ANKA recipient line, Bergreen, which constitutively expresses GFP under the control of the *HSP70* promoter (Kooij et al., 2012) (Figure 2 B: clone 2; Figure S1 A: clone 1).

At this stage, we could already conclude that the successful generation of clonal *uis12(-)*-knockout lines demonstrates that the gene is not essential for normal progression of asexual blood stages, although *UIS12* is expressed considerably in schizonts (Figure 1 B).

### Impaired gametocyte development of *uis12(-)* parasites

After successful selection of *uis12(-)* parasites, we first examined asexual and sexual blood stage growth. To this end, C57BL/6 mice (*n*=5) were infected by intravenous injection of either 10,000 *uis12(-)* or WT-infected erythrocytes. Starting three days after infection the mice were monitored by daily microscopic examination of thin blood films, and parasitemia and gametocytemia were quantified (Figure 2 C and D). Parasitemia increased daily in wild-type- and *uis12(-)*-infected mice to a similar extent. On day 7, wild-type and *uis12(-)*-infected mice started developing symptoms indicative of onset of experimental cerebral malaria, supporting the notion that *uis12(-)* parasites display virulence typically seen in infections with *P. berghei* ANKA parasites.

The blood films were also analyzed for the presence of male and female gametocytes (Figure 2 D). This analysis revealed that, in contrast to normal asexual blood infection, gametocytemia was significantly reduced in *uis12(-)* parasites as compared to the wild-type. *Uis12(-)* gametocyte numbers were already slightly reduced on day four after infection, when the first fully mature gametocytes became visible. The difference between WT and *uis12(-)* parasites increased further over time. Examination of size and morphology of male or female gametocytes and ookinetes revealed no difference between the two parasite lines (Figure 2 E). The presence of ookinetes indicates that *uis12(-)* gametocytes are able to fertilize and subsequently form a zygote that develops into an ookinete. Both, male and female gametocytes were found in the expected female biased proportions. Together, the data indicate that UIS12 is indispensable for asexual growth, but plays an important role for gametocytogenesis. Analysis of the second *uis12(-)* clone corroborated these findings (Figure S1 B and C).

### Exflagellation is markedly reduced in *uis12(-)* parasites

The potential of male gametocytes to produce motile microgametes is an active process and termed exflagellation. In both *uis12(-)* clonal lines, exflagellation events were detected very rarely by microscopic examination of undiluted blood from *uis12(-)* infected mice at any given time point. Yet, this already showed that *uis12(-)* male gametocytes are able to produce motile male gametes, albeit at very low frequencies. To quantify the exflagellation centers, C57BL/6 mice were injected intravenously with a high dose of 10^7^ blood stages. Four days later, parasitemia, gametocytemia and the number of exflagellation centers were enumerated (Figure 2 F, *uis12(-)* clone 1). Interestingly, in this experiment, the parasitemia was slightly, but significantly, higher in *uis12(-)*-infected mice. In agreement with our first experiment (Figure 2 D), gametocytemia was 8.6-fold reduced in *uis12(-)* parasite-infected mice. Exflagellation was not visible under these conditions, *i*.*e*. 1:25 diluted blood, in *uis12(-)*-infected mice (Figure 2 F). Even after an extended incubation or when examined the following days, exflagellation was not detectable. In conclusion, male *uis12(-)* gametocytes retain the ability to progress to functional microgametes, but at a very low rate. The defect in exflagellation is more severe than the reduction in gametocytemia indicating additive defects in absence of *UIS12* upon progression from blood infection to transmission to the insect vector.

### Reduced oocyst formation by *uis12(-)* gametocytes, but failure to produce sporozoites

The severe defect in *in vitro* exflagellation implied a large impact in mosquito colonization. However, after transmission to *Anopheles stephensi* mosquitos, *uis12(-)* parasites were able to establish midgut infections and produce oocysts, showing that early events of midgut colonization are not abolished *in vivo* (Figure 3). The mean mosquito infection prevalences were 54% (±18%) for wild-type (*n*=5) and 46% (± 23%) for *uis12(-)* (*n*=5).

**Figure 3:**
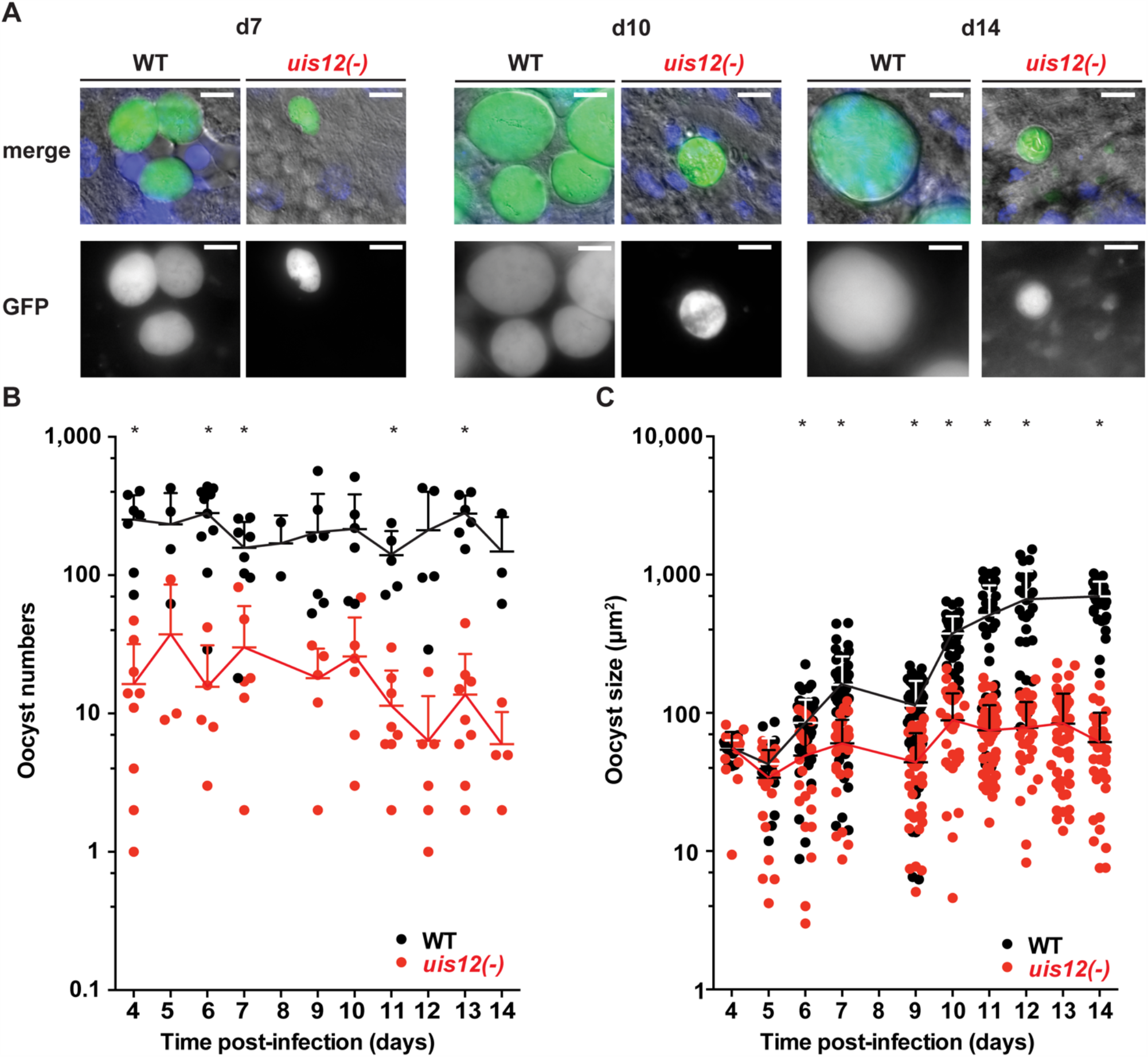
Disruption of *UIS12* leads to a reduced oocyst size and a defect in sporozoite formation. (A) Shown are representative live micrographs of *uis12(-)* and wild-type oocysts 7, 10 and 14 days after an infectious blood meal. GFP (green) is constitutively expressed under the *HSP70* promoter (*P. berghei* ANKA Bergreen)(Kooij et al., 2012). *Uis12(-)* clone 2 was used for this experiment. Nuclei were stained with Hoechst (blue). Note the reduced density and size of *uis12(-)* oocysts, and absence of sporozoites in *uis12(-)* oocysts. Magnification, 630-fold; scale bars, 10 µm. (B, C) Time course of oocyst numbers per infected mosquito midgut (B) and oocyst size (µm^2^) (C) at different time points after midgut infection (*uis12(-)* clone 2, red; WT, black). *, *p*<0.01 (multiple t-tests).

This finding prompted us to quantify oocyst maturation. To this end, midguts were isolated daily starting on day 4 after mosquito infection up until day 14. Oocyst numbers and morphology as well as their ability to produce sporozoites were analyzed by fluorescence microscopy (Figure 3, *uis12(-)* clone 2). In *uis12(-)-*infected mosquitoes, the number of oocysts per midgut was more than 10-fold reduced as compared to wild-type, consistent with reduced gametocyte numbers and the very low exflagellation rate (Figure 3 B). Strikingly, in the absence of *UIS12* oocysts were markedly smaller in size (Figure 3 A and C). While up until day 4 the oocyst size was comparable between both groups, over time *uis12(-)* oocysts displayed an abnormal morphology. Their size remained unchanged, whereas the WT oocyst diameter increased ∼10-fold over the following ten days. This finding indicates that the early events of oocyst development, including transformation of the motile ookinete to the spherical oocyst, and the initial increase in biomass (Carter et al., 2007) are reduced, but not completely abolished. An additional defect leads to arrested oocyst maturation. The analysis of the second *uis12(-)* clone corroborated these findings (Figure S1 D-F).

Upon close examination at later time points, *uis12(-)* oocysts displayed little or no nuclear staining, harbored many large vacuoles, and frequently were surrounded by an apparently wrinkled oocyst wall (Figure 3 A, Figure S1 F). As a consequence, no sporozoites were seen inside *uis12(-)* oocysts. This lack of sporozoite formation was confirmed by analysis of extracted midguts and salivary glands; no midgut- or salivary gland -associated sporozoites were detectable. In WT-infected midguts sporoblast formation was observed inside the oocysts starting around day 11 after infection, and from day 12 onwards oocysts started to fill with sporozoites (Matuschewski, 2006) (Figure 3 A). In contrast, *uis12(-)* midguts remained free of sporozoites even until late time points of mosquito infection, excluding delayed sporozoite formation.

To obtain independent support for the complete absence of sporozoites in *uis12(-)*-infected mosquitoes, we inoculated C57BL/6 mice with salivary gland extracts of 20 *uis12(-)*-infected mosquitoes. In addition, we exposed C57BL/6 mice to ∼100 *uis12(-)*-infected mosquitoes (Table 1, both clones). In sharp contrast to control infections with wild-type-infected mosquitoes under similar conditions, none of the attempts to transmit *uis12(-)* parasites resulted in blood stage infections (Table 1). This defect in transmission confirms that *uis12(-)* parasites are unable to complete their development inside the mosquito.

**Table 1:** No malaria transmission by *uis12(-)*-infected mosquitoes.

| Parasites | Route of injection<sup>a</sup> | Infected/ Injected | Pre-patency (days)<sup>b</sup> |
| --- | --- | --- | --- |
| WT 507cl | 10,000 sporozoites i.v. | 5/5 | 3 |
| *uis12(-)* clone 1 | mosquito bite (~100 mosquitoes) | 0/5 | N.A. |
|  | salivary gland extract i.v. (20 mosquitoes) | 0/4 | N.A. |
| WT Bergreen | mosquito bite (~100 mosquitoes) | 3/3 | 3 |
| *uis12(-)* clone 2 | mosquito bite (~100 mosquitoes) | 0/3 | N.A. |
|  | salivary gland extract i.v. (20 mosquitoes) | 0/3 | N.A. |

<sup>a</sup> C57BL/6 mice were exposed to the bites of ~100 infected mosquitoes for 15-20 minutes or injected intravenously (i.v.) with salivary gland extract isolated from 20 mosquitoes (*uis12(-)*) and, as a control, 10,000 sporozoites (WT). Data are pooled from two separate experiments.
<sup>b</sup> the pre-patent period represents the days until detection of the first infected erythrocytes by microscopic thin blood film examination after sporozoite inoculation. n.a., not applicable.

### Deregulation of transcripts important for mosquito transmission in *uis12(-)* blood stages

We hypothesized that due to the presence of the two RRM motifs, UIS12 is likely involved in post-transcriptional control of gene expression. Reduced gametocyte production, exflagellation and oocyst development detected in *uis12(-)* parasites suggests that UIS12 might be regulating target mRNAs and alter expression of the respective proteins, which in turn play important roles in these stages.

To profile the transcriptome of *uis12(-)* parasites, we performed a comparative microarray analysis on *uis12(-)* and wild-type mixed blood stage RNAs (Figure 4, Figure S2). Since the first defect is detected during gametocyte formation, we limited our analysis to mixed blood stages, that include the asexual blood stages, from which gametocytes originate. Attempts to retrieve sufficient *uis12(-)* gametocytes for a microarray failed. The microarray analysis was performed with two biological replicates (R1 and R2), both with *uis12(-)* clone 1. For both replicates, parasitemia of the samples were comparable (R1: WT 6.2%; *uis12(-)* 6.3%; R2: WT 6.4%; *uis12(-)* 8.7%). As expected, gametocytemia was reduced in *uis12(-)* parasites (R1: WT 0.2%; *uis12(-)* 0.06%). Each microarray experiment was performed by dual-color hybridizations, and a color-swap dye-reversal was included in order to compensate for dye-specific effects. In this microarray analysis, the -fold change of the expression levels of 2,890 *P. berghei* genes could be successfully compared (Figure 4 A, Figure S2 A, D).

**Figure 4:**
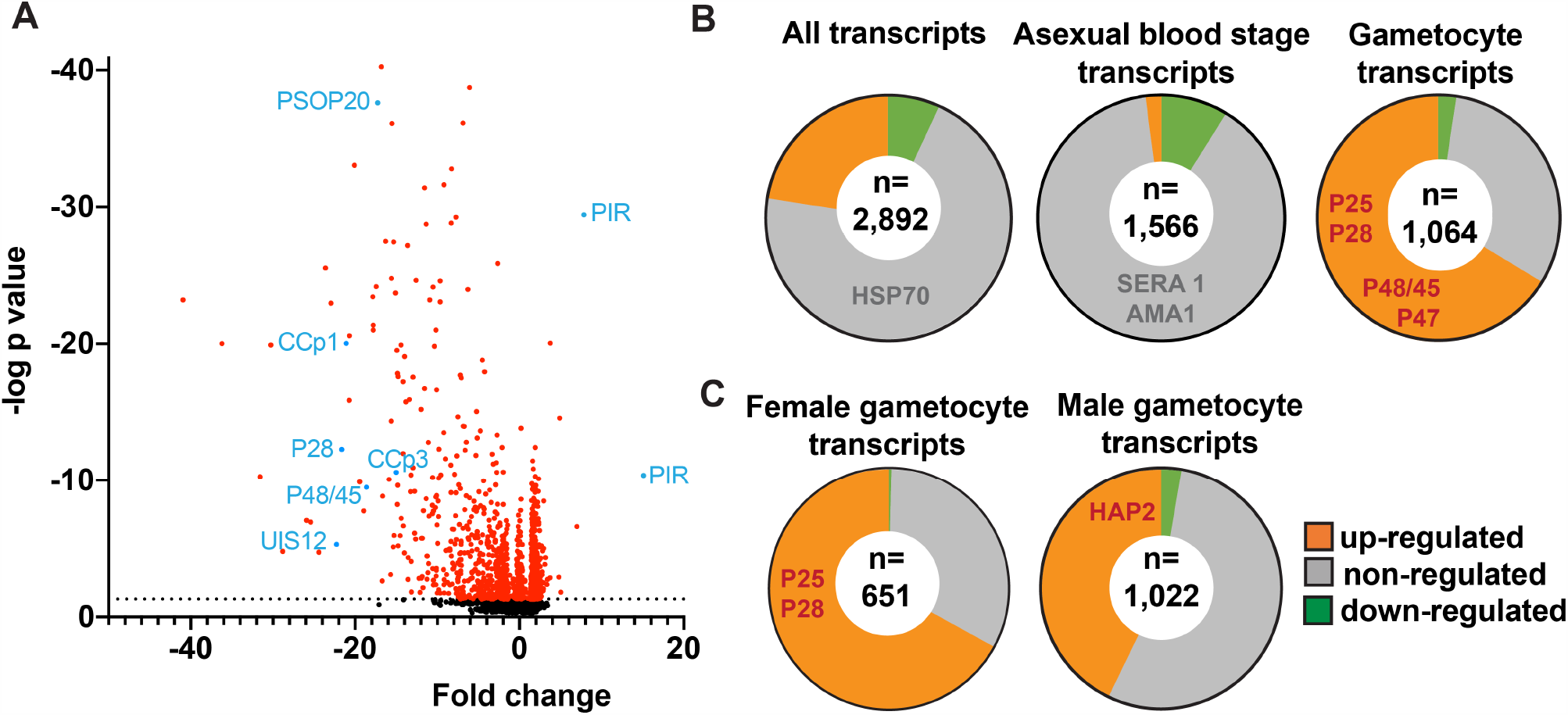
Down-regulation of distinct mRNAs coding for signature gamete-, ookinete- and oocyst-specific proteins in *uis12(-)* blood stage parasites. Shown are mean values of microarray results of total RNA isolated from mixed infected erythrocytes from *uis12(-)* (clone 1)*-* and WT-infected mice from two biological replicates. (A) Shown is a volcano-plot illustrating the fold change of the expression levels and the negative log *p*-values of all analyzed 2,890 *P. berghei* genes. The dotted black line represents a *p*-value of 0.05 and all transcripts with a *P*-value <0.05 are shown in red. Exemplary transcripts are highlighted and labeled in blue. (B) Pie charts displaying the proportions of up- (green >2), non- (grey), and down- (orange <-2) regulated transcripts amongst all transcripts (upper left), blood stage-specific transcripts (center) and gametocyte-specific transcripts (upper right) (Otto et al., 2014). Exemplary transcripts are listed in the respective region of the chart. The number of transcripts analyzed is shown in a white circle inside the center. (C) Pie charts displaying the proportions of up- (green >2), non- (grey), and down- (orange <-2) regulated transcripts amongst female (left) and male (right) gametocyte-specific transcripts (Yeoh et al., 2017). The number of transcripts analyzed is shown in a white circle inside the center.

Overall, absence of *PbUIS12* correlated with a global perturbation of 30% of all mRNA expression levels (threshold, <-2 or >2). More transcripts were down-regulated (646, 23%) than up-regulated (199, 7%). The mean down-regulation was -6.4-fold, whereas the mean up-regulation was +2.5-fold (GEO Series accession number: GSE152686). As expected, the *UIS12*-knockout was independently confirmed by the microarray analysis; *UIS12* transcripts were down-regulated by - 22.3-fold (Table 2). Within the 25 most down-regulated transcripts known proteins required for the switch to the mosquito vector, CCp1 (LAP2), LAP5, P48/45, and P28, as well as numerous previously unrecognized proteins are represented (Table 2). The list of the 25 most up-regulated genes contains, among others, PIR proteins (Table S1). Remarkably, many transcripts that were down-regulated more than 10-fold in *uis12(-)* blood stages, are reported to be expressed in gametocytes (Hall et al., 2005). We also analyzed the proportions of up-, down- and non-regulated transcripts and found distinctions between stage-specific RNAs, as defined by RNA sequencing data (Otto et al., 2014). For instance, gametocyte transcripts were defined by >2-fold up-regulation in gametocytes as compared to asexual blood stages (Figure 4 B, Table 3, Figure S2 B, E). In good agreement with normal replication rates, very few asexual blood stage-specific transcripts were down-regulated in *uis12(-)* parasites. Instead, the proportion of up-regulated transcripts was highest for asexual blood stage-specific mRNAs, which might reflect over-representation of these stages due to defects in gametocytogenesis. However, signature transcripts, like *AMA1* (−0.3), *HSP70* (−1.6), *MSP1* (1.7) and *MSP8* (1.5), were neither up-nor down-regulated in the absence of *UIS12* (Figure 4 A, Table 3). We also noted that the mRNA levels of many signature mosquito stage-specific proteins were down-regulated in *uis12(-)* mixed blood stages. In addition, we also compared the proportions of up-, down- and non-regulated transcripts and found distinctions between male- and female-specific transcripts, as defined by RNA sequencing data (Yeoh et al., 2017) (Figure 4 C, Figure S2 C, F). While 67% of the female gametocyte transcripts were down-regulated in absence of *UIS12*, only 43% of male-specific transcripts were down-regulated, indicating differential impact of UIS12 on male and female gametocyte-specific transcripts in mixed blood stages.

**Table 2:** The top 25 down-regulated genes in blood stages of *uis12(-) vs*. wild-type.

| Gene ID | Product description | Mean -fold change R1<sup>a</sup> R2<sup>b</sup> | -fold change R1<sup>a</sup> | -fold change R2<sup>b</sup> |
| --- | --- | --- | --- | --- |
| PBANKA_0806800 | unknown | -41.0 | -37.5 | -44.4 |
| PBANKA_1449000 | candidate microgamete surface protein MiGS | -36.2 | -25.4 | -47.0 |
| PBANKA_1349400 | candidate DNA replication complex GINS protein | -31.6 | -29.0 | -34.2 |
| PBANKA_1340400 | candidate lactate dehydrogenase | -30.2 | -24.2 | -36.3 |
| PBANKA_1400400 | fam-b-protein | -28.8 | -14.3 | -43.4 |
| PBANKA_1361600 | E1-E2 ATPase | -25.9 | -25.9 | -25.9 |
| PBANKA_1040300 | fam-b-protein | -25.4 | -18.5 | -32.3 |
| PBANKA_1315300 | LCCL domain-containing protein (LAP5) | -24.4 | -27.4 | -21.4 |
| PBANKA_0413000 | unknown | -23.6 | -29.3 | -18.0 |
| PBANKA_1246600 | cleft-like protein 1 | -22.9 | -9.7 | -36.2 |
| PBANKA_0506200 | RNA binding protein (UIS12) | -22.3 | -31.7 | -12.8 |
| PBANKA_0514900 | ookinete surface protein P28 | -21.6 | -15.5 | -27.8 |
| PBANKA_1300700 | LCCL domain-containing protein (CCp1) | -21.1 | -20.7 | -21.4 |
| PBANKA_0719700 | unknown | -20.7 | -19.2 | -22.2 |
| PBANKA_0707100 | inner membrane complex protein 1i | -20.7 | -26.5 | -15.0 |
| PBANKA_1334900 | unknown | -20.4 | -20.1 | -20.7 |
| PBANKA_1353800 | unknown | -20.1 | -26.1 | -14.1 |
| PBANKA_1432400 | perforin-like protein 2 | -19.4 | -16.9 | -21.9 |
| PBANKA_0608600 | candidate EGF-domain protein | -19.0 | -23.9 | -14.0 |
| PBANKA_1400900 | unknown | -18.9 | -20.3 | -17.5 |
| PBANKA_1359600 | 6-cysteine protein (P48/45) | -18.6 | -21.0 | -16.2 |
| PBANKA_1106000 | unknown | -17.9 | -22.2 | -13.5 |
| PBANKA_1414800 | unknown | -17.9 | -19.6 | -16.1 |
| PBANKA_1120400 | candidate inner membrane complex protein 1j | -17.8 | -18.9 | -16.7 |
| PBANKA_1318600 | unknown | -17.8 | -15.9 | -19.7 |

<sup>a</sup> biological replicate 1<sup>b</sup> biological replicate 2

**Table 3:** Examples for deregulation of signature vector-stage transcript in *uis12(-)* parasites.

|  | Known blood stage transcripts |  | Known transmission stage transcripts |  |
| --- | --- | --- | --- | --- |
| Up-regulated | Pb235 | 2.8 |  |  |
| Unaffected | AMA1 | -0.3 | AP-2 G* | -0.9 |
|  | MSP8 | 1.5 | p230p | -0.3 |
|  | HSP70 | -1.6 | NPT1 * | -1.3 |
|  | SERA3 | 1.7 | Puf2 | -1.9 |
|  | MSP1 | 1.7 |  |  |
| Down-regulated<br>($< 10$-fold) | | | CITH* | -2.3 |
|  |  |  | Actin II | -2.8 |
|  |  |  | CDPK4 | -3.1 |
|  |  |  | MAPK 2 | -3.3 |
|  |  |  | GAP45 | -5.1 |
|  |  |  | AP2-0 | -5.7 |
|  |  |  | CDPK3 | -5.8 |
|  |  |  | Plasmepsin VI | -6.3 |
|  |  |  | HAP2 | -7.5 |
|  |  |  | MDV1 | -10 |
| Down-regulated<br>($> 10$-fold) | | | CCp5* | -11.2 |
|  |  |  | CCp4* | -12.2 |
|  |  |  | CCp3* | -15 |
|  |  |  | P47 | -15 |
|  |  |  | CCp2* | -15.3 |
|  |  |  | P25 | -17.1 |
|  |  |  | P48/45 | -18.6 |
|  |  |  | CCp1 * | -21.1 |
|  |  |  | P28 | -21.6 |

Shown are exemplary blood and transmission stage transcripts. The mean -fold change represents steady-state mRNA levels in *uis12(-)* parasites compared to the corresponding WT parasite stages. \*, genes with a knock-out phenotype similar to *uis12(-)*.

Next, we wanted to independently confirm the microarray results by quantitative RT-PCR of selected transcripts on blood stage and gametocyte mRNA from *uis12(-)* and WT samples (Figure 5). This analysis revealed a good agreement with the mean -fold changes determined in the microarray for gametocyte- and blood stage-specific transcripts. The gametocyte-expressed transcripts *Puf1, DOZI, UIS1/IK2, SET* and *Actin II* (Deligianni et al., 2011; Gomes-Santos et al., 2011; Mair et al., 2006; Muller et al., 2011; Pace et al., 2006) showed similar -fold changes in both assays. Similarly, the blood stage-expressed transcripts *AMA1* and *MSP1* remained un-regulated. For some transcripts, *e*.*g*. male development factor (*MDV1*) (Lal et al., 2009), the reduction in *uis12(-)* parasites was even lower by quantitative RT-PCR in comparison to the microarray data. qRT-PCR on gametocyte mRNA samples were consistent with the data from asexual blood stages (Figure 5, *uis12(-)* clone 1). Overall, we could confirm the results of the microarray on these samples, indicating that the down-regulation of the gametocyte transcripts is a direct result of a specific UIS12-dependent regulation and not merely a consequence of the absence of gametocytes in *uis12(-)* mixed blood stages.

**Figure 5:**
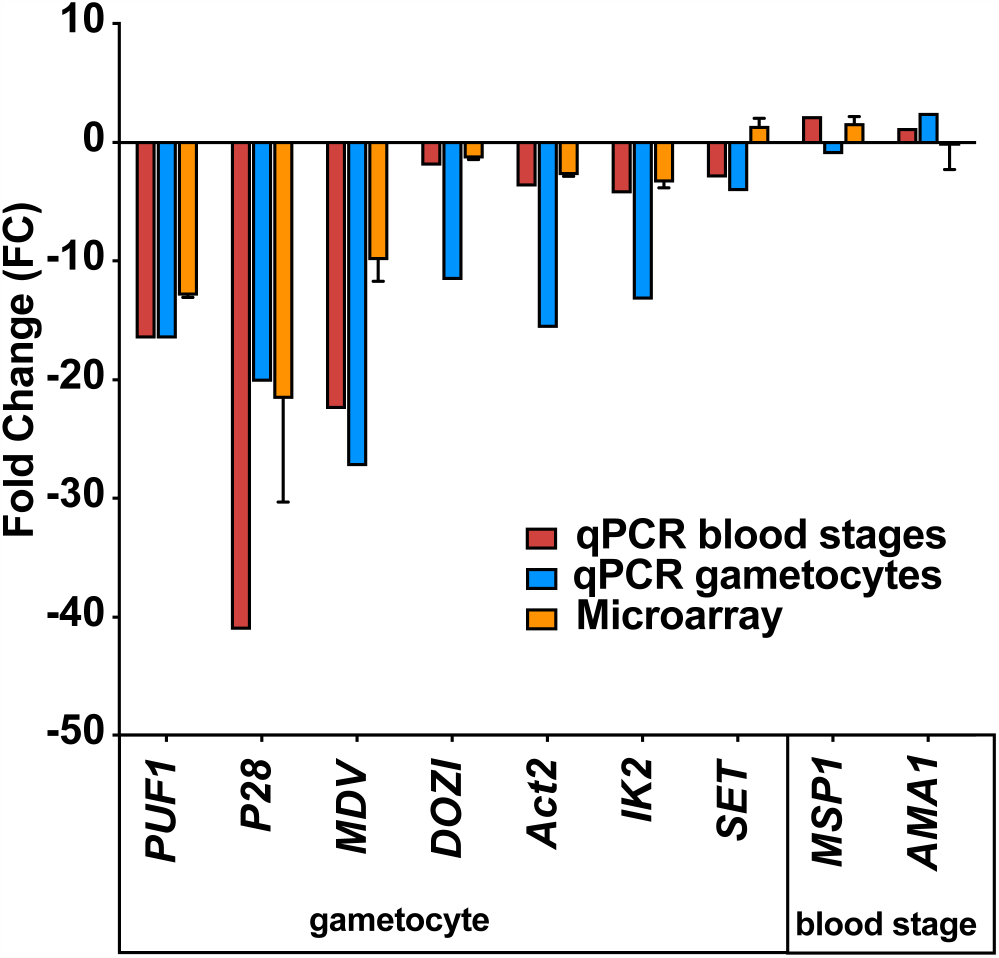
Confirmation of down-regulation of selected mRNAs by quantitative RT-PCR. Quantitative RT-PCR on selected blood stage and gametocyte transcripts validate the results of the microarray analysis. Shown are -fold changes of steady state mRNA levels of *uis12(-)* clone 1 compared to wild-type mixed blood stages for selected genes (*Puf1, P28, MDV, DOZI, Actin II, UIS1/IK2, SET, MSP1*, and *AMA1*). qRT-PCR data were normalized to the steady state levels of *HSP70* mRNA. The microarray data represent mean values of biological replicate 1 and 2 ±SD.

### Gene ontology analysis of the deregulated transcripts in *uis12(-)* parasites

We next performed a gene ontology (GO) enrichment analysis (Ashburner et al., 2000) (Tables S2 and S3). According to this analysis the 199 up-regulated and the 646 down-regulated transcripts are involved in 21 and 42 different biological processes, as described by distinct GO terms, respectively. Examples for GO terms of the set of down-regulated genes include reproduction, cell communication and phosphorylation (Table S2), in good agreement with the known roles of kinases in gametogenesis (Invergo et al., 2017). Examples for GO terms of the set of up-regulated genes include RNA processing and histone acetylation (Table S3), further indicative of perturbations in gene expression and mRNA translation in absence of *UIS12*.

### A 10-nucleotide motif shared by up-regulated transcripts in *uis12(-)* parasites

Since UIS12 contains two RNA recognition motifs and the combination of at least two RRM domains allows the continuous recognition of a 8-10 nucleotide-long sequence motif with high affinity (Maris et al., 2005), we screened all ORFs, and 1,000 base pairs of the flanking 5’ UTR and 3’ UTR sequences of genes up-regulated >2.5-fold in replicate 1 for common 8-10 nucleotide-long motifs using MEME (Bailey et al., 2009) using the respective portions of the non-regulated transcripts as reference. While this search did not return a motif in either the 5’ or 3’ UTR sequences, we found a 10 nucleotide-long motif (U/C-U_/A_-U-C/U-U/_A_-U-U/C-U-U-C) in the ORF (Figure 6 A). This signature was found 751 times in 174 of the 220 up-regulated sequences (e-value: 4.2×10^−4^). We restricted our search to upregulated transcripts, since down-regulation can at least be partially attributed to fewer gametocytes in mixed *uis12(-)* blood stages.

**Figure 6:**
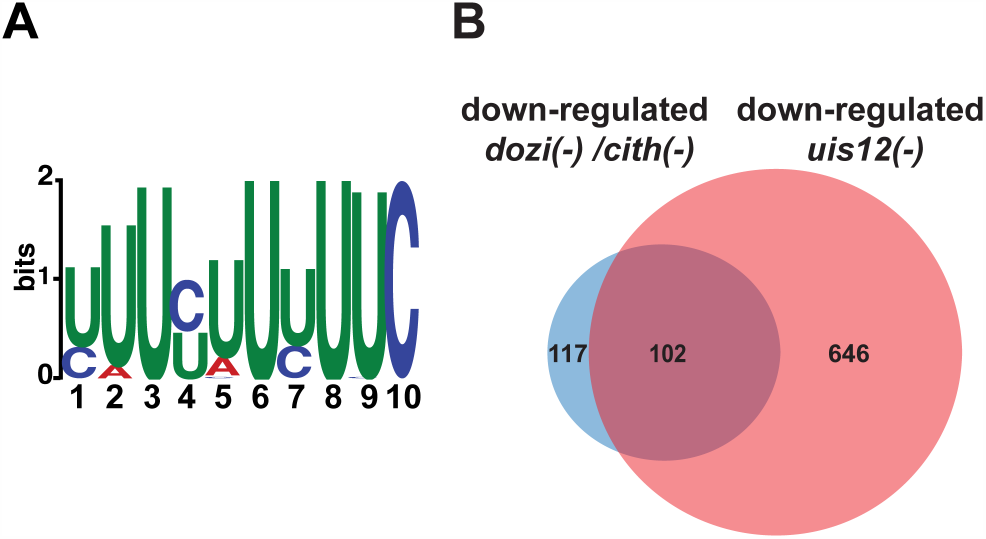
A motif is shared by up-regulated transcripts in *uis12(-)* parasites, and the comparison of *dozi(-), cith(-)* and *uis12(-)* target transcripts reveals possibly overlapping functions. (A) Shown is a graphic display of the 10-nucleotide U-rich motif found in the ORF of transcripts that are up-regulated in absence of *UIS12* in the microarray with biological replicate 1. The size of the depicted nucleotide represents its probability at the respective position. (B) Venn diagram comparing shared down-regulated transcripts of the gametocyte microarray analyses performed for *dozi(-)* and *cith(-)* with a -fold change lower than -1 (Mair *et al*., 2010) and down-regulated transcripts in mixed blood stages of *uis12(-)* with a mean -fold change lower than -2.

Together, this 10 nucleotide-long U-rich motif might represent a signature of UIS12 interaction with target mRNAs during blood infection, which is apparently required for proper preparation for host switching.

## Discussion

In this study, deletion of the candidate RNA binding protein UIS12 led to the discovery of a potential post-transcriptional regulator of gametocytogenesis and mosquito transmission. Our results underscore the key roles of a fine-tuned transcriptional control and post-transcriptional regulation during transmission of the malaria parasite between hosts. When *UIS12* is absent, fewer gametocytes are produced and *in vitro* exflagellation is severely impaired, yet not abolished. The defects of *uis12(-)* parasites during host switching cannot be compensated for in subsequent steps in the *Plasmodium* life cycle, and *uis12(-)* infection leads to fewer oocysts, which fail to undergo oocyst maturation. We can correlate these defects with a set of signature gametocyte and mosquito-stage transcripts that are down-regulated in *uis12(-)* parasites already during blood infection. Parasites lacking *UIS12* produce ∼80 % fewer gametocytes. Both male and female gametocytes account for this reduction, indicative of a role of *UIS12* in early events of gametocytogenesis, perhaps in conjunction with transcriptional control by AP2-G, AP2-I, AP2-FG and AP2-G2. Gene deletion of *AP2-G* resulted in a complete loss of gametocytes and the deletion of *AP2-G2* in a major reduction of gametocytes (Sinha et al., 2014). While *AP2-G* expression was unaffected in *uis12(-)* parasites, *AP2-G2* was not included in our microarray screen.

In contrast to a broad role in gene regulation, such as translational inhibition by phosphorylation of eIF2α, RRM-containing RNA binding proteins are mainly known for their tight RNA-specific control of RNA processing, export, and stability. A typical RNA recognition motif consists of 80 amino acids that folds into an αβ sandwich and adopts a β_1_α_1_β_2_β_3_α_2_β_4_ topology. In this conformation the motif is tailored to bind to single-stranded RNA with 4 conserved aromatic residues in the β_1_ and β_3_ sheet, which largely mediate RNA binding. The presence of tandem RRM domains ensures high sequence specificity (Maris et al., 2005). *Plasmodium* falciparum possesses at least 189 candidate RBP-proteins, which belong to 13 families and comprise 3.5% of all annotated genes. The RRM domain is represented with 8 families and 72 proteins in *P. falciparum*, and often multiple RRM domains are present in a protein. In good agreement with the complex regulation network required for stage conversion, 27% of all RBPs show elevated expression during the gametocyte stage (Reddy et al., 2015).

Whether UIS12 is a positive regulator of gametocytogenesis and how it influences this process awaits further molecular and biochemical analysis, including RNA-pull down studies to capture UIS12 target transcripts. The role of UIS12 in promoting gametocytogenesis shares similarities with another RNA-binding protein, the pumilio-familiy RNA-binding protein *Puf1* which was shown to function in *P. falciparum* gametocyte maturation, and especially of female gametocytes (Shrestha et al., 2016). Both proteins likely inhibit asexual propagation and, thereby, promote gametocyte formation. A related pumilio-familiy RNA-binding protein, *Puf2*, exerts the corresponding function in sporozoites and prepares them for transmission to the vertebrate host (Gomes-Santos et al., 2011; Miao et al., 2010; Muller et al., 2011). We note that UIS12 likely executes a second function, similar to the one analyzed in the present study, in sporozoites, the stage where it was originally identified (Matuschewski et al., 2002), but the complete block in sporogony prevented us from studying this by classical reverse genetics. It, thus, still remains unclear if *UIS12* also plays a role in the host switch back to the mammalian host. Of note, in sporozoites translational repression of *UIS12* was described, and the protein is only detectable in liver stages (Lindner et al., 2019). Accordingly, a stage-specific knockout, which depletes *UIS12* during sporogony is required to study its cellular roles for sporozoite transmission and liver infection.

The few remaining *uis12(-)* gametocytes were able to form male and female gametes, fertilize, and form ookinetes. Accordingly, numbers of oocysts are extremely low in *uis12(-)-*infected *Anopheles* mosquitoes. Complete life cycle arrest in *uis12(-)* infections occurs at an early time-point after oocysts are formed, indicating that the events leading to oocyst formation are not impaired (Carter et al., 2007). Some oocysts appeared empty, and other oocysts shrunken and forming wrinkles. This defect is reminiscent of the wrinkled walls of oocysts during and after sporozoite release, previously shown in scanning electron microscopy studies (Chen et al., 1984; Meis et al., 1992).

Our microarray analysis revealed a sharp drop in the steady state mRNA levels of numerous well-characterized *Plasmodium* genes. The *P. berghei* LCCL protein family members CCp1 and lectin adhesive-like protein 5 (LAP5) are expressed in female gametocytes and ookinetes. *Pb*LAP proteins are important for sporogenesis inside oocysts, yet known to be expressed already in gametocytes (Lavazec et al., 2009; Pradel et al., 2004; Raine et al., 2007; Saeed et al., 2010; Trueman et al., 2004). The surface proteins of the 6-cysteine repeat protein family P48/45 are essential for male gametocyte fertility (van Dijk et al., 2001). P25 and P28 are surface proteins of sexual stages, with distinct roles in ookinete to oocyst transformation (Siden-Kiamos et al., 2000, Tomas et al,.

2001). Interestingly, some transcripts that are severely depleted, including P28, AP2-O, CCP2 and CCP4, have been reported to be translationally controlled (Lasonder et al., 2016; Saeed et al., 2013; Vervenne et al., 1994). Strikingly, the function of some of these repressed targets extends far beyond the ookinete stage, and the corresponding proteins are important for oocyst and sporozoite formation, illustrating the complexity of regulation of posttranscriptional control of gene expression in gametocytes (Lasonder et al., 2016). A deregulation of repressed mRNAs in absence of *UIS12* might partially explain the observed inability to complete oocyst development. However, the abundant *UIS12* mRNA levels in ookinetes suggest a potential additional checkpoint of post-transcriptional control by *UIS12*, that is around the time, when transcription of mosquito stage-specific target genes is orchestrated by a family of closely related Apetala-2 proteins, AP2-O 1-4, in ookinetes (Kaneko et al., 2015; Modrzynska et al., 2017). And this additional checkpoint in ookinetes might be the reason for the pleiotropic effects observed upon absence of *UIS12*, finally resulting in the aberrant maturation of oocysts. Due to the low *UIS12* transcript levels in oocysts a direct role at this point in the life cycle is less likely. We, thus, propose that UIS12 exerts multiple regulatory roles at consecutive life cycle stages, and, hence, is together with Puf2 one of the first known *Plasmodium* RNA binding proteins with pleiotropic functions for life cycle progression (Gomes-Santos et al., 2011; Lindner et al., 2013; Miao et al., 2013; Miao et al., 2010; Muller et al., 2011).

The GO-term enrichment of down-regulated transcripts in *uis12(-)* shows perturbation of many kinases and GO-term enrichment of up-regulated transcripts revealed perturbation of gene expression. A MEME search in the ORFs, 5’UTRs and 3’UTRs of the up-regulated transcripts identified in the microarray did reveal a 10-nucleotide U-rich motif in the ORFs unique to the analyzed subsets when compared to sequences that were not perturbated in absence of *UIS12*. We wish to point out that down-regulation of gametocyte signature transcripts in *uis12(-)* blood stages can be largely attributed to the at least 3.5-fold reduction in gametocyte numbers, yet the levels of down-regulation ranged from a low (−2, e.g. *Puf2*) to more than 15-fold (e.g. *CCP1* and *CCP2*). By qPCR we could confirm the down-regulation of some of the gametocyte specific transcripts such as P28 in a gametocyte enriched mRNA preparation of *uis12(-)* blood stage parasites, indicating that the down-regulation of gametocyte transcripts might not only be a consequence of the absence of gametocytes in *uis12(-)* mixed blood stages, but a direct result of a specific UIS12-dependent regulation. Further analyses employing RNA-Seq or single cell transcriptomics on synchronized and purified stages, ranging from ring stages to ookinetes, are warranted to gain an in-depth understanding of the mRNA repertoire that is under UIS12 control. Since the shared signature motif was identified in up-regulated transcripts, this would argue against a role of UIS12 as an mRNA stabilizing factor. At least two alternative scenarios emerge; one potential, albeit unusual, mechanism of UIS12 could be binding and destabilizing of transcripts. The reverse, and perhaps more likely, function would be an inhibitory role towards a partner protein, which in turn stabilizes target mRNAs. According to such a scenario, up-regulation of transcripts in the absence of UIS12 might be a consequence of increased mRNA stability. We consider the identification of a nucleotide motif a good starting point for future work, and are hesitant to propose a molecular mechanism at this early stage of discovery.

In gametocytes, the conserved DOZI/CITH RNA storage complex (Guerreiro et al., 2014; Mair et al., 2006; Mair et al., 2010) is central to translational repression. For instance, the DOZI/CITH complex is involved in translationally repressing *P25, P28, PlasmepsinVI, AP2*-O and *GAP45*. These transcripts are also contained on our list of down-regulated transcripts in *uis12(-)* blood stages. Thus, we were curious how many targets overlap when we compared the reported 117 genes down-regulated in both *dozi(-)* and *cith(-)* as determined by microarray on gametocytes (Mair et al., 2010) with the 646 down-regulated transcripts in *uis12(-)* blood stages (<-2-fold change), we found a surprisingly high overlap, of 102 of the 117 transcripts (Figure 6 B). The RNA-helicase DOZI was previously shown to be a key regulator of *P25* and *P28* transcript stability, as well as of 731 other transcripts, and they all share a conserved U-rich nucleotide motif mainly in the 5’ and 3’ UTR (Braks et al., 2008; Guerreiro et al., 2014; Hall et al., 2005). The abundance of U residues in the DOZI and UIS12 motifs could hint at a related mechanism. The key differences are that the U-rich motif in our study is found in the ORFs of target genes, while the U-rich motif bound by the DOZI complex is located in 5’ and 3’ regions of its targets (Braks et al., 2007). Also, the U-rich motif identified for DOZI binding has a length of 47 nucleotides, whereas the motif identified herein is only 10 nucleotides long. Despite these clear distinctions a potential functional link of these two proteins remains an attractive possibility, since UIS12 and DOZI have >100 overlapping down-regulated targets (Figure 6 B).

We note that, in marked contrast to *UIS12*, gene deletion of *DOZI* leads to a developmental arrest of the fertilized female gamete, a process that is not abolished in the absence of UIS12. In all three knockout lines, *uis12(-), dozi(-)*, and *puf2(-)* parasites the mRNA levels of *P25* and *P28* were perturbated. While in *dozi(-)* and *uis12(-)* the transcripts were down-regulated, in *puf2(-)* an upregulation was described, indicating involvement in distinct shared and opposing regulatory processes. On the other hand, PlasmepsinVI is translationally repressed by *Pf*Puf2 (Miao et al., 2013) and reduced in *uis12(-)* parasites, similar to the meiotic recombinase *DMC1* (Mlambo et al., 2012). Both knockouts as well as the *CCP* knockouts resemble the defects of *uis12(-)*. Clearly, further work is warranted to decipher the molecular mechanisms that orchestrate the sequential post-transcriptional control events and contributions of the regulatory molecules.

In conclusion, our experimental genetics analysis of the *Plasmodium berghei* candidate RNA binding protein *UIS12* uncovered broad perturbations of mRNA expression during blood infection, including upregulation of several hundred transcripts, which share a U-rich 10 nucleotide-long motif in their open reading frames. The observed mRNA deregulation correlates with pleiotropic defects after blood stage propagation during sexual stage development, mosquito midgut colonization and oocyst maturation, ultimately leading to a complete block in the *Plasmodium* life cycle inside the *Anopheles* vector.

## Methods

### Ethics statement

All animal work conducted in this study was carried out in strict accordance with the German ‘Tierschutzgesetz in der Fassung vom 18. Mai 2006 (BGBl. I S. 1207)’, which implements the directive 86/609/EEC from the European Union and the European Convention for the protection of vertebrate animals used for experimental and other scientific purposes. The protocol was approved by the ethics committee of MPI-IB and the Berlin state authorities (LAGeSo Reg# G0469/09, G0294/15).

### Parasites and experimental animals

*P. berghei* ANKA cl507 parasites that constitutively express GFP under the PbEF1α promoter and *P. berghei* ANKA Bergreen parasites that express the green fluorescent protein GFP under the PbHSP70 promoter throughout their life cycle have been used in our experiments (Franke-Fayard et al., 2004; Kooij et al., 2012).

All mouse experiments were performed with female NMRI or C57BL/6 mice purchased from Charles River Laboratories (Sulzfeld, Germany) or bred in house. NMRI mice were used for transfection experiments, blood stage infections and transmission to *Anopheles stephensi* mosquitos. C57BL/6 mice were used for sporozoite and blood stage infections, exflagellation essays and ookinete cultures.

### Gene deletion of *P. berghei UIS12*

The *P. berghei UIS12* gene deletion was performed by double crossover homologous recombination of a replacement pB3D-knockout plasmid (van Dijk et al., 1995). The gene knockout construct targeting the endogenous *UIS12* locus comprised a 647 bp *UIS12* 5’ UTR and a 524 bp *UIS12* 3’ UTR sequence flanking a *T. gondii DHFR*/TS pyrimethamine-resistance cassette. Table S4 lists the oligonucleotides used for generation of the knockout construct. Prior to transfection, plasmids were linearized by restriction digest. A digest with *Sac*II and *Kpn*I produced a fragment consisting of the pyrimethamine resistance cassette flanked by the *UIS12* 5’ and 3’ UTRs. Transfection by electroporation of cultured schizonts was performed as previously described (Janse et al., 2006b). Two independent parasite lines, one in *P. berghei* Bergreen and one in *P. berghei* cl507 recipient lines were generated and phenotypically characterized. For genotyping of recombinant parasites, knockout- and wild-type-specific PCR fragments were amplified from genomic DNA of parental and clonal lines with the oligonucleotides listed in Table S4.

### Quantitative real-time PCR (qRT-PCR)

Quantitative real-time PCR was used to quantify *UIS12* gene expression and for comparing gene expression in *uis12(-)* and wild-type (WT) mixed blood stages and gametocytes. For mixed blood stage analyses, C57BL/6 mice were infected i.v. with 10,000 iRBC, and 7 days after infection whole blood was isolated. The gametocyte samples were generated by infecting phenylhydrazine-treated NMRI mice with 10^6^ iRBC. Three to four days later, when gametocytemia reached the highest levels, 12.5 mg sulfadiazine was supplemented to 1 liter of drinking water. Within 48 h the sulfadiazine treatment eliminated the asexual stages, which was confirmed by a Giemsa-stained thin blood film. Next, the infected blood was harvested. To isolate mixed blood stages or gametocytes, whole blood was cleaned from white blood cells and erythrocytes using cellulose-glass bead columns and saponin (0.3%; Sigma-Aldrich) lysis. Parasites were stored in 1ml TRIzol (Invitrogen) or RLT-Buffer with β-mercaptoethanol (Qiagen). Parasite RNA was isolated with the RNeasy Kit (Qiagen) or TRIzol-Reagent (Sigma-Aldrich). DNase treatment with Quiagen RNase free DNase Set or Ambion Turbo DNA free Kit was performed to remove residual genomic DNA.

Reverse transcription to cDNA was done using the RETROscript Kit (Ambion). The qPCR was performed on parasite cDNA that was generated from total RNA of selected *P. berghei* life cycle stages. Quantitative real-time PCR was performed as previously described (Silvie et al., 2008) using Power SYBR Green PCR Master Mix and the StepOnePlus Real-Time PCR System (both Applied Biosystems). All procedures were performed according to the manufacturer’s instructions. Life cycle expression data were normalized to GFP that is constitutively expressed under the *EF1α* promoter in *P. berghei* 507cl. Mixed blood stage expression data for comparison of knockout and WT parasites were normalized to *PbHSP70*. The relative transcript abundance was determined by the 2^ΔΔCT^ method. For oligonucleotide sequences refer to Table S4.

### *Plasmodium* life cycle: Parasite asexual and sexual growth, mosquito infection and sporozoite transmission

For asexual blood stage and gametocyte growth curves, C57BL/6 mice were injected intravenously with 10,000 infected red blood cells. From day 3 to 7, Giemsa (VWR)-stained thin blood films were microscopically examined to determine parasitemia and gametocytemia. Parasitemia and gametocytemia are defined as the percentage of red blood cells infected with asexual stages or gametocytes, respectively.

To analyze male exflagellation, C57BL/6 mice were intravenously injected with 10^7^ infected red blood cells (iRBC). After four days, parasitemia and gametocytemia were determined. Next, five microliters of tail blood were diluted 1:25 in ookinete medium (RPMI 1640, 10% FCS and 50mM xanthurenic acid, pH 8.0) and incubated for 12 minutes at room temperature. For the next six minutes exflagellation centers were counted with help of a Neubauer chamber.

Ookinete cultures were set up, as previously described (Weiss et al., 1997). Four days after the intravenous injection of 10^6^ iRBC into C57BL/6 mice pre-treated one day earlier with phenylhydrazine 97% (Sigma-Aldrich, 6.18µl/ml) 1 ml of fresh mouse blood was mixed with 10 ml of ookinete medium. After 20 hours incubation at 20°C, at 80% humidity the culture was examined for development of ookinetes and ookinetes were purified with anti-P28-coupled magnetic beads.

*Anopheles stephensi* mosquitoes were kept at 28°C (non-infected) or 20°C (infected) under a 14 h light/ 10 h dark cycle at 80% humidity. Four days after the transfer of 10^7^ iRBC, when exflagellation reached a maximum, mosquitoes were infected as described (Vanderberg, 1975). After 4 to 14 days, midguts were isolated and oocyst development analyzed by fluorescent microscopy (Zeiss AxioObserver Z1). For visualization of nuclei, HOECHST 33342 (Invitrogen) was added at a 1:1,000 dilution. Images were analyzed using the FIJI processing (oocyst diameter) and/ or Photoshop (oocyst numbers) software.

Midguts and salivary glands were analyzed at day 14-18 and day 18-30, respectively, for presence of sporozoites.

To test *Plasmodium* transmission, age-matched female C57BL/6 mice were either subjected to the bites of ∼100 infected mosquitos or inoculated with 10,000 sporozoites, or for *uis12(-)* the salivary gland extract of 20 mosquitos, in RPMI intravenously. After three days Giemsa-stained thin blood films were analyzed for presence of infected red blood cells.

### Microarray expression profiling of *P. berghei* blood stages

The microarray was performed on two biological replicate samples of *uis12(-)* and wild-type with comparable parasitemia. One replicate was generated by infecting one C57BL/6 mouse each with 10,000 *uis12(-)* or wild-type blood stages intravenously (R1, biological replicate 1). After 7 days of infection mixed blood stages were isolated from whole blood. For both samples, the parasitemia was comparable (*uis12(-)* 6.3%, WT 6.2%). The other replicate (R2, biological replicate 2) was generated by injecting one NMRI mouse with 10^7^ *uis12(-)* or wild-type blood stages and mixed blood stage parasites were harvested after 3 days (*uis12(-)* 8.7%, WT 6.4%).

Whole blood was isolated and cleaned from white blood cells and erythrocytes using cellulose-glass bead columns and saponin (0.3%; Sigma-Aldrich) lysis. About 10^8^ parasites were stored in 1ml TRIzol (Invitrogen). Total RNA was isolated using glycogen as a carrier according to the ‘TRIzol Reagent RNA preparation method’ (Invitrogen). The amount of RNA was determined by OD 260/280 nm measurement using a ‘NanoDrop 1000’ (peQlab) spectrophotometer. The RNA size, integrity and the amount of total RNA were measured with a ‘Bioanalyzer 2100’ using an ‘RNA Nano 6000 Microfluidics Kit’ (both Agilent genomics). RNA labeling was performed with the two color ‘Quick Amp Labeling Kit’ (Agilent genomics) using a 1:5 mixture of ‘FullSpectrum MultiStart Primer’ (Systembio). mRNA was reverse transcribed and amplified using the primer mixture. The RNA was split and labeled with Cyanine 3-CTP and Cyanine 5-CTP, respectively. After precipitation, purification and quantification, labeled samples were hybridized to the Agilent ‘4×44K custom-commercial microarrays’ according to the manufacturers protocol. Microarray experiments were performed as dual-color hybridizations. A color-swap dye-reversal was performed in order to compensate specific effects of the dyes and to ensure statistically relevant data analysis (Churchill, 2002; Wernersson et al., 2007). Scanning of the microarrays was performed with 5 µm resolution and the extended mode using a ‘High Resolution Microarray Laser Scanner’ (G2505, Agilent Technologies). Raw microarray image data were extracted and analyzed with the ‘G2567AA Image Analysis / Feature Extraction software’ (Version A.10.5.1.1, Agilent Technologies). On reporter and sequence level, the extracted MAGE-ML files were further analyzed with the ‘Rosetta Resolver Biosoftware’ (7.2.2.0 SP1.31). All subsequent data analysis was performed with the Microsoft Excel, Prism and the PlasmoDB (http://plasmodb.org/plasmo/) database. *P. berghei* microarrays were designed on the OligoWiz (http://www.cbs.dtu.dk/services/OligoWiz/) server that included annotated open reading frames (ORF) from the whole genome of *P. berghei*. ORF specific probes were designed as sense oligonucleotides of the coding strand (Wernersson et al., 2007). The aimed oligonucleotide length was between 50 and 60 bases. To avoid self-hit, the maximal homology was 97 %, the cross hybridization maximum length was at 80 % and random primed position preference scores within OligoWiz were chosen. Depending on these parameters and the sequence input length, each ORF was covered by different numbers of specific oligonucleotide probes. In total 41,133 specific probe sequences were uploaded to eArray (Agilent Technologies, https://earray.chem.agilent.com/earray/) as expression application with a customer specified feature layout.

The microarray data discussed in this publication have been deposited in NCBI’s Gene Expression Omnibus (Edgar et al., 2002) and are accessible through GEO Series accession number GSE152686.

### Statistical analysis and online bioinformatic tools

Statistical significance was determined with the help of Graph Pad Prism software, using Mann-Whitney test for comparison of parasitemia, gametocytemia and exflagellation dot-plots and multiple t-tests (one per row) for all growth curves.

The identification and positioning of the two RRM domains of the UIS12 orthologous proteins has been performed by NCBI WEB batch-CD Search Tool (Marchler-Bauer et al., 2015). Protein comparison was done by NCBI protein blast (blastp) with default settings, no compositional adjustment and without low complexity filter. Sequence coverage of all *Plasmodium* orthologues was higher than 90%.

Gene ontology (GO) enrichment analysis (Ashburner et al., 2000) was performed with help of the Gene Ontology enrichment analysis tool from PlasmoDB, with the default settings and the redundant GO terms were removed with help of the REVIGO software.

To screen for common 8-10 nucleotide-long motifs we used the MEME webtool (Bailey et al., 2009), screened all ORFs, 5’UTR sequences and 3’UTR sequences (1,000 nucleotides per gene for ORFs) of transcripts up-regulated >2.5-fold and compared them to the respective gene parts of the non-regulated (−1.5 to −1.5) transcripts from our microarray analysis of biological replicate 1.

## Author Contributions

K.Ma, O.S. and K.Mü designed the experiments; K.Mü performed experiments and analyzed data; H-J.M. performed and analyzed the microarray; K.Ma and K.Mü wrote the paper; all authors commented and revised the manuscript.

## Data Availability Statement

The datasets generated and analyzed for this study can be found in the NCBI’s Gene Expression Omnibus (Edgar et al., 2002) and are accessible through GEO Series accession number GSE152686.

